# When “Time Online” Tells Us Little: Duration of Social Media Use does not Influence Sleep Disruption or Psychopathology in Adolescents

**DOI:** 10.64898/2026.10.04.756572

**Authors:** Morgan Lott, Sophia Koesterer, Eleanor Mackellar, Amanda McCleery, Gerta Bardhoshi, Jonathan Platt, Bengi Baran

## Abstract

**Background:** Adolescent mental health problems have risen alongside increasing sleep disruption and widespread social media use (SMU). However, the mechanisms linking these factors remain unclear. This study concurrently examined objectively measured sleep and SMU in relation to psychopathology to test whether SMU moderates the sleep–mental health association and to characterize day-to-day bidirectional relations among sleep, affect, mood, and SMU during the school year.

**Methods:** Eighty-five adolescents (ages 12–18) completed 14 days of wrist actigraphy, smartphone-based ecological momentary assessments of mood and affect, and objective SMU duration tracking. Cross-sectional models tested whether SMU moderated associations between sleep and psychopathology, and multilevel daily models examined within-person lagged associations among sleep, SMU, and affect, controlling for age and sex. False discovery rate correction was applied.

**Results:** Shorter average sleep duration and greater variability in sleep duration between weekdays and weekends were associated with higher psychopathology symptoms, primarily externalizing problems. SMU duration neither moderated nor directly predicted psychopathology or sleep. Within-person analyses showed that longer-than-usual sleep predicted more positive and less negative affect and lower next-day SMU, though these associations did not survive correction for multiple comparisons.

**Conclusions:** Sleep disruption showed consistent associations with adolescent psychopathology, whereas the amount of time spent on social media did not. These findings suggest that duration-based measures of SMU may incompletely capture adolescents’ digital experiences and highlight the need for more granular, context-sensitive indicators of online behavior in understanding youth mental health.

**Key points:** *What’s known:*

- Adolescent mental health problems commonly co-occur with sleep disruption and high social media use, yet evidence for a mechanistic link between social media use and mental health remains inconsistent.

*What’s new:*

- This study integrates actigraphy, objective social media use data and daily ecological momentary assessments to examine real-world behavior over two weeks. Greater variability in sleep duration, but not time spent on social media, was associated with psychopathology symptoms.

*What’s relevant:*

- Findings highlight sleep as a potentially modifiable target for improving adolescent emotional well-being and suggest that time-based social media restrictions alone may have limited clinical, educational or policy impact.

## INTRODUCTION

Several reports point to an alarming increase in the prevalence of mental health conditions among adolescents over the past two decades(Sappenfield et al., 2018, Bitsko et al., 2022) highlighting the need to identify modifiable developmental risk processes. In parallel, multiple large-scale studies have documented a decline in adolescent sleep duration, indicating worsening sleep health at the population level(Twenge et al., 2017, Matricciani et al., 2012). These concerning trends in adolescent mental health and sleep have coincided with the widespread availability of smartphones and the rapid expansion of social media platforms. By 2022, 95% of U.S. teens reported having access to a smartphone, and nearly half said they engage with social media “almost constantly”(Vogels et al., 2022). While this pervasive digital engagement has led researchers and policymakers to call for precautionary measures and stronger regulation of social media platforms, evidence for a mechanistic relationship between social media use (SMU) and youth mental health is lacking(Hartwell et al., 2024). Sleep is fundamental for brain development, cognitive function, and emotion regulation, and meta-analytic evidence shows bidirectional associations between sleep disturbances and symptoms of depression, anxiety, and poorer well-being (Bacaro et al., 2024). Yet, a largely untested possibility is that the relationship between sleep disruption and psychopathology may vary as a function of SMU. Social media use may interfere with sleep through time displacement, engagement in the hours immediately preceding sleep, heightened emotional arousal, or increased blue light exposure such that the same degree of sleep disruption may be associated with greater psychopathology among adolescents with higher levels of digital engagement. To test this proposed mechanistic account, we objectively measured sleep via actigraphy and SMU duration via smartphone-based data over 14 consecutive days during the school year, allowing us to capture real-world behavior with high temporal precision. We tested whether higher SMU moderated the relationship between sleep quality and psychopathology and evaluated daily bidirectional associations among sleep, SMU, affect, and mood.

Puberty is a unique period of rapid maturational changes in brain circuitry governing sleep-wake regulation. Adolescent sleep is characterized by a slower buildup of sleep pressure(Jenni and Carskadon, 2004) and a circadian phase delay(Roenneberg et al., 2004) that promotes a biological preference for later bedtimes and wake times. Increasing academic and social demands and extracurricular activities further delay bedtimes, while early school start times remain unchanged or even shift earlier during high school. This convergence of biological, psychosocial, and structural factors, described as the “*Perfect Storm”*(Carskadon, 2011), may lead to chronically insufficient and misaligned sleep in youth(Owens, 2014, Foundation, 2006, Pallesen et al., 2011, Urner et al., 2009). In turn, inadequate or misaligned sleep is known to contribute to a range of adverse outcomes including the onset or worsening of psychiatric symptoms, cognitive deficits, academic problems, increased risk of motor vehicle accidents, suicidal thoughts and behaviors, and metabolic disturbances such as obesity(Crowley et al., 2018, Carskadon et al., 2004). Daily fluctuations in sleep and mood also appear to be linked. Several studies have found that disrupted sleep predicts higher next-day negative affect and anxiety (Fuligni and Hardway, 2006, Ben Simon et al., 2020), although the strength and direction of these associations vary across studies(Shen et al., 2022, Kouros et al., 2022, Xie et al., 2024). Individual and contextual factors further moderate these associations, with stronger relations between sleep and affect observed among youth with higher externalizing symptoms and those with greater nighttime screen use(Dong et al., 2024, Evans et al., 2024, Carlo et al., 2015).

Associations between SMU and adolescent mental health have been widely examined, although meta-analytic estimates generally indicate modest correlations (r =.10 - .15) (Bozzola et al., 2022, McCrory et al., 2020, Vidal et al., 2020, Verduyn et al., 2017, Damodar et al., 2021, Keles et al., 2019, McCrae et al., 2017, Memon et al., 2018, Radtke et al., 2022, Sedgwick et al., 2019, Valkenburg et al., 2022, Hancock et al., 2022, Ivie et al., 2020, Appel et al., 2020, Meier and Reinecke, 2020). Such small effects suggest that SMU is not uniformly associated with mental health risk across adolescents. Instead, its relevance to mental health may depend on co-occurring processes, such as disrupted sleep. One possibility is that greater SMU amplifies the association between sleep disruption and psychopathology, such that adolescents experiencing both greater sleep disruption and higher SMU may show greater symptom burden than would be expected from either factor alone. For instance, SMU may delay bedtime, displace sleep opportunities and contribute to difficulties in maintaining sleep (Scott et al., 2019). Sleep disruption appears especially pronounced if social media engagement occurs immediately before sleep (Levenson et al., 2017, Lemola et al., 2015, Nagata et al., 2023). The emotional and social content of online interactions may contribute to a state of hyperarousal, making it more difficult to initiate sleep(Cain and Gradisar, 2010, Scott and Woods, 2018). In addition, exposure to artificial light from backlit screens suppresses melatonin production in a dose-dependent manner and may disrupt circadian timing (Tähkämö et al., 2019, West et al., 2011). This may potentially exacerbate the biologically-driven tendency toward delayed sleep timing during adolescence. Collectively, these findings raise the possibility that SMU may not only be associated with sleep disruption itself but may also exacerbate the emotional consequences of poor sleep, thereby strengthening associations between sleep disturbance and psychopathology. Complementing these pathways, evidence also suggests that the relationship may be bidirectional, with sleep disruption potentially contributing to greater technology use (Bauducco et al., 2024). Adolescents having trouble initiating sleep may use technology to occupy periods of wakefulness or potentially as a strategy for regulating affect during the sleep-onset period(Daniels et al., 2023). This possibility underscores the importance of examining temporal associations in both directions rather than assuming that greater technology use precedes sleep disruption.

However, to our knowledge, no prior study has concurrently measured sleep, SMU and mental health using objective, high-frequency and longitudinal methods in youth, an approach critical for testing mechanistic hypotheses and identifying clinically meaningful risk profiles. Although one recent study combined self-reported bedtime SMU, actigraphy, and daily affect assessments in young adults, bedtime SMU was not associated with poorer sleep or next-day affect (Das-Friebel et al., 2020). Whether similar patterns emerge in adolescents, who engage with social media more frequently and across a broader range of daily contexts, remains unclear. Further, prior research has relied heavily on cross-sectional designs or self-report measures, leaving unclear whether daily variations in objectively measured sleep and digital behavior contribute to fluctuations in mood and affect. To address these gaps, we combined actigraphy-based sleep assessment, objective SMU data, and ecological momentary assessments of affect over a two-week period, enabling analysis of both between-person differences and within-person daily dynamics. At the between-person level, we hypothesized that sleep disruption would be positively associated with psychopathology symptoms and that, consistent with the moderation hypothesis described above, this association would be stronger among adolescents with greater social media engagement. At the within-person level, we examined a distinct question: whether day-to-day fluctuations in sleep and SMU were temporally associated with changes in daily affect, providing novel insight into short-term temporal dynamics.

## METHODS

### Participants

Participants were 85 children and young adults aged 12-18 (mean: 16.3±2.1 yrs) recruited via electronic flyers at local public middle and high schools or via introductory psychology courses at the University of Iowa. Inclusion criteria were (1) fluency in English for both the participant and their primary caregiver (2) no self-reported history of intellectual and developmental disabilities or neurological disorders; (3) no history of seizures or unexplained loss of consciousness; (4) no history of head injury resulting in loss of consciousness or neurological sequalae, and (5) no travel across time zones or shift work in the past two weeks. All study procedures were approved by the Institutional Review Board at the University of Iowa. After the study procedures were explained, written informed consent was obtained from participants aged 18 years, and assent was obtained from participants under 18 years along with consent from their primary caregiver. Participants < 18 yrs received monetary compensation for their participation, while college freshmen (aged 18) received course credit.

### Procedures

#### Study Design

The study included two in-person laboratory visits separated by two weeks of remote data collection. During the initial visit, participants (and caregivers for those under age 18) completed the computerized Kiddie Schedule for Affective Disorders and Schizophrenia (KSADS; (Townsend et al., 2020) that included pre-interview self/caregiver report measures and structured interviews administered by trained research staff to characterize the sample with respect to present and lifetime diagnoses of psychiatric conditions based on DSM-5 criteria. Participants also completed self-report measures on sleep and mental health (described below), received actigraphy devices, and were trained on smartphone-based ecological momentary assessments (EMA). At Visit 2, actigraphy devices were returned and SMU logs were obtained directly from smartphone tracking tools, minimizing recall bias.

#### Objective Sleep Measurement

Participants wore an ActTrust 2 actigraphy device (Condor Instruments, São Paulo, Brazil) on their non-dominant wrist for two weeks to continuously record activity, ambient light, and body temperature. Devices sampled at 25 Hz in 30-s epochs, and data were analyzed in ActStudio (Condor Instruments, São Paulo, Brazil) using the proportional integration mode (PIM). Participants pressed the device’s event marker button at bedtime and upon waking. Sleep scoring was conducted using a semi-automatic procedure consistent with published actigraphy scoring recommendations (Ancoli-Israel et al., 2015). Actigraphy records were initially scored with the ActStudio algorithm and subsequently reviewed by trained scorers. Event-marker data and EMA sleep diary entries were used to verify and refine algorithm-derived estimates of nightly lights-off and lights-on times. When participant-reported information was missing, incomplete, or discrepant, scorers used consensus review of activity and ambient light signals to determine the most likely lights-off and lights-on times, with activity patterns weighing most heavily in scoring decisions. Primary measures included daily bedtime and wake up times, sleep onset latency (SOL; minutes from lights-off to sleep onset), total sleep time (TST), wake after sleep onset (WASO; minutes awake after sleep onset), and sleep efficiency (SE; percentage of time asleep while in bed). Weekday-weekend differences in midsleep timing (social jetlag) and total sleep duration (sleep debt) were also calculated to index sleep regularity.

#### Ecological Momentary Assessments

Participants completed EMA of mood, affect, and sleep using the Avicenna Research smartphone application (Toronto, Canada; formerly *Ethica*). The app delivered two daily surveys (see Supplementary Tables S1-2). Morning surveys (available 6 am-noon) included sleep diary items on bedtime, wake time, and sleep onset latency. Evening surveys (available 7pm-2am) assessed mood and affect using a modified 12-item version (Shen et al., 2022) of the Positive and Negative Affect Schedule (PANAS-X (Watson, 1999)) rated on a 5-point Likert scale (1 = very slightly or not at all; 5 = extremely). Daily positive and negative affect scores were calculated from the evening EMA responses by averaging the relevant PANAS-X items. Mood elation was measured with a statement rated from 0 (do not agree at all) to 100 (completely agree): “Right now I feel much more giddy, silly, or happy than usual.”

#### Social Media Usage

Daily social media use was quantified via the smartphone native tracking tools (Apple Screen Time or Android Digital Wellbeing), which automatically log time spent on social media applications. Both systems automatically categorize applications into predefined usage categories (e.g., social networking/social media) and quantify foreground screen engagement while excluding passive background activity. Participants used either Apple (n = 75) or Android (n = 10) devices and provided consent to share daily usage data with the research team over the two-week monitoring period.

#### Self-Report Measures

The Achenbach System of Empirically Based Assessments (ASEBA) Youth Self-Report (YSR)(Achenbach, 1999) was used to assess the presence and severity of mental health problems. Raw counts were converted to age- and sex-normed t-scores, which provide standardized continuous indices of internalizing and externalizing psychopathology severity. Daytime sleepiness was measured with the Pediatric Daytime Sleepiness Scale (PDSS(Drake et al., 2003)). Puberty was assessed using a self-administered Pubertal Development Scale(Carskadon and Acebo, 1993), and home environment structure was evaluated with the Confusion, Hubbub, and Order Scale (CHAOS(Matheny et al., 1995)), with higher scores indicating environmental confusion.

### Statistical Analyses

Analyses were conducted in IBM SPSS Statistics (v29) and in R (v4.4.1) within RStudio (v2024.04.1+748) to examine relationships among sleep, mental health and SMU. Two analytic goals were pursued. First, to characterize <u>between-person associations</u> of sleep, SMU and psychopathology we averaged the actigraphy-based sleep metrics across the two-week observation period to derive a stable and objective estimate of each participant’s typical sleep quality. We then tested whether this overall sleep quality was associated with psychopathology symptoms, and whether SMU duration moderated this association. Second, <u>within-person daily associations</u> of sleep, SMU and affect were examined using multilevel models with days nested within participants. Lagged models evaluated temporal associations between daytime indices and overnight sleep. Specifically, models tested whether daily SMU or affect predicted sleep that night, and whether nightly sleep predicted affect and SMU the following day. Statistical significance was set at *p* < .05 (two-tailed). False discovery rate (FDR) correction(Benjamini and Hochberg, 1995) was applied to control for multiple comparisons.

#### Cross-Sectional Analyses: Associations Between Sleep, Social Media Use, and Psychopathology

Moderation analyses examined whether SMU altered the strength of the relationship between sleep and psychopathology symptoms. Models were conducted using the PROCESS Macro(Hayes and Little, 2018) (v5.0) for SPSS. SMU was operationalized as the standardized average daily duration of use across the two-week period. Mental health problems were indexed by ASEBA YSR Total Problems t-scores. Sleep measures were derived from actigraphy and reduced using principal component analysis (PCA) to minimize redundancy among daily metrics. SMU was specified as the moderator, with age and sex included as covariates. Predictors and moderators were mean-centered prior to computing interaction terms, and heteroscedasticity-consistent standard errors were used. Conditional effects of sleep on mental health were probed at low (−1 SD), mean, and high (+1 SD) levels of SMU, Johnson-Neyman regions of significance were examined and multiple comparisons were FDR corrected.

#### Daily-Level Analyses

*Temporal Dynamics of Sleep, Affect, and Social Media Use* Predictors and outcomes were temporally aligned for each participant, with lags applied when appropriate (e.g., sleep on day *t* predicting affect or SMU on day *t + 1*; affect or SMU on day *t* predicting sleep that night). For analyses requiring a lag, the first observation was excluded because lagged values were undefined. All time-varying predictors were person-mean centered to isolate within-person effects, and participant-level means across the monitoring period were entered simultaneously to estimate between-subject effects. Thus, within-person terms indexed whether deviations from an adolescent’s own average predicted daily outcomes, whereas participant-level means indexed stable between-subject differences in average sleep, SMU, and affect across the monitoring period.

Linear mixed-effects models with a random intercept for participant accounted for repeated measures over the study period. Each model included age and sex as covariates. Daily fluctuations in sleep were indexed using actigraphy-derived TST and WASO. Daily positive affect, negative affect and mood elation were derived from the evening EMA surveys. Daily SMU reflected total minutes spent on social media apps. Within-person associations tested (1) sleep predicting next-day affect and mood, (2) sleep predicting next-day SMU, (3) affect or mood predicting same-night sleep, (4) SMU predicting same-night sleep, (5) affect or mood predicting same-day SMU, and (6) SMU predicting same-day affect or mood. The same models simultaneously estimated between-subject associations using participant-level means of the time-varying predictors. Analyses were conducted in R using the lme4(Bates et al., 2007)and lmerTest(Kuznetsova et al., 2017) packages. FDR correction was applied to control for multiple comparisons across both the within-person and between-person effects.

## RESULTS

### Participant Characteristics

Table 1 shows participant demographic characteristics. In this community sample, daytime sleepiness scores on the PDSS ranged from 0 to 29, and 33% of participants scored above the proposed cutoff indicating excessive daytime sleepiness(Meyer et al., 2018). Average daily social media use also showed substantial variability, ranging from 0 to more than 6.5 hours per day (mean= 182.4 ± 100.2 mins). Actigraphy data were available for 71 participants (data were not obtained from 14 participants, 6 male and 8 female, due to device malfunction or temporary unavailability). Actigraphy-derived sleep patterns are shown in Table 2 and revealed significantly reduced sleep duration on weekdays compared with weekends, whereas daily SMU duration did not differ between weekdays and weekends. Fifteen participants met DSM-5 criteria for a current disorder during the research-based KSADS interview (See supplementary Table S3 for the list of these research diagnoses). These participants did not differ from the rest of the sample with respect to age (F(1,83)=.02, *p*=.88), overall time spent on social media (F(1,83)=2.12, *p*=.15), average actigraphy measured sleep duration (F(1,83)=.82, *p*=.37) or sleep efficiency (F(1,83)=.22, *p*=.64) but reported higher daytime sleepiness (F(1,83)=7.48, *p*=.008).

**Table 1.**
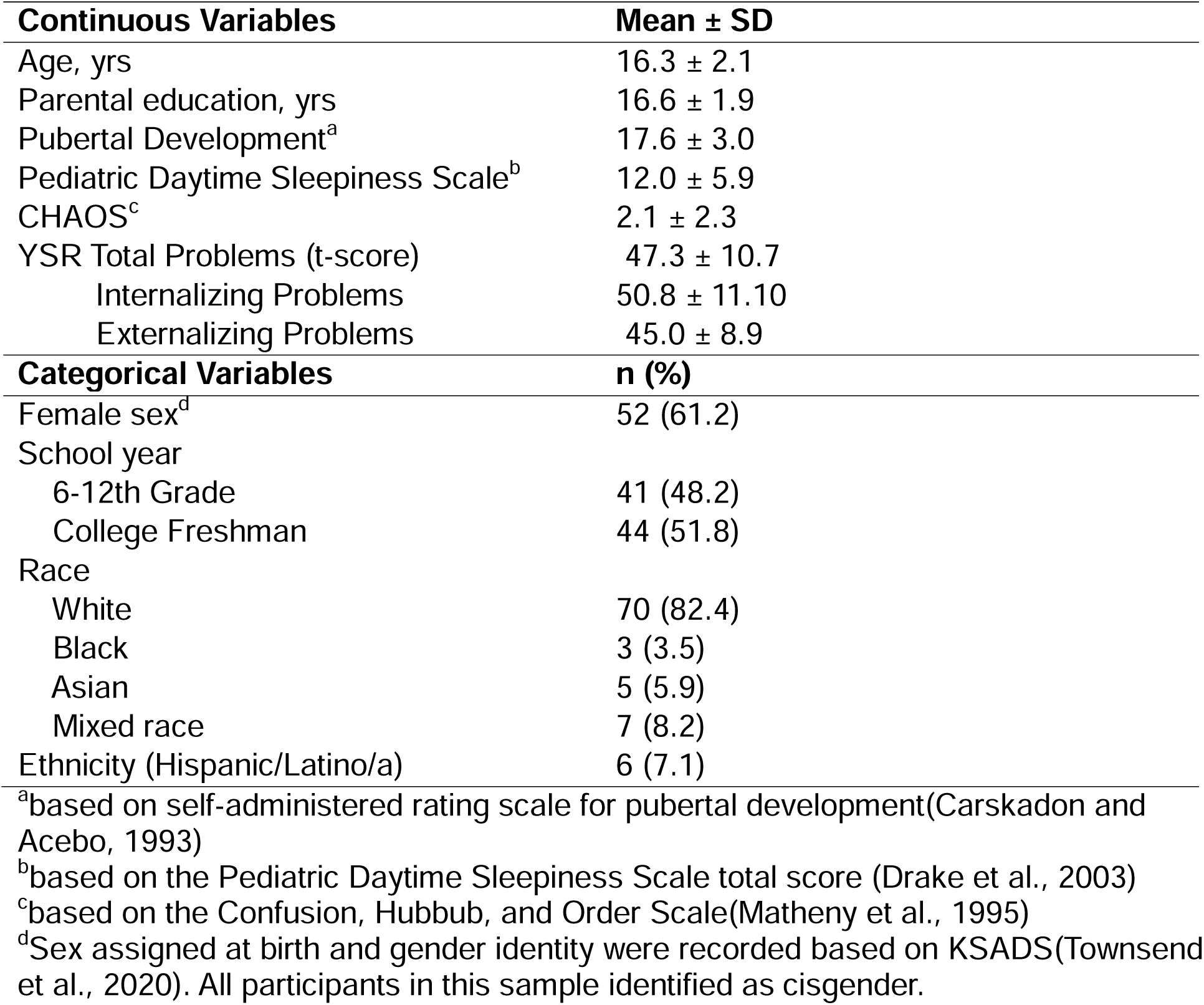
Demographic Characteristics of the Sample (N = 85)

| **Continuous Variables** | **Mean ± SD** |
| --- | --- |
| Age, yrs | 16.3 ± 2.1 |
| Parental education, yrs | 16.6 ± 1.9 |
| Pubertal Development<sup>a</sup> | 17.6 ± 3.0 |
| Pediatric Daytime Sleepiness Scale<sup>b</sup> | 12.0 ± 5.9 |
| CHAOS<sup>c</sup> | 2.1 ± 2.3 |
| YSR Total Problems (t-score) | 47.3 ± 10.7 |
| Internalizing Problems | 50.8 ± 11.10 |
| Externalizing Problems | 45.0 ± 8.9 |
| **Categorical Variables** | **n (%)** |
| Female sex<sup>d</sup> | 52 (61.2) |
| School year |  |
| 6-12th Grade | 41 (48.2) |
| College Freshman | 44 (51.8) |
| Race |  |
| White | 70 (82.4) |
| Black | 3 (3.5) |
| Asian | 5 (5.9) |
| Mixed race | 7 (8.2) |
| Ethnicity (Hispanic/Latino/a) | 6 (7.1) |

<sup>a</sup>based on self-administered rating scale for pubertal development(Carskadon and Acebo, 1993)
<sup>b</sup>based on the Pediatric Daytime Sleepiness Scale total score (Drake et al., 2003)
<sup>c</sup>based on the Confusion, Hubbub, and Order Scale(Matheny et al., 1995)
<sup>d</sup>Sex assigned at birth and gender identity were recorded based on KSADS(Townsend et al., 2020). All participants in this sample identified as cisgender.

**Table 2.** Descriptive statistics for objective measures of sleep and social media use averaged across the two-week monitoring period separated by weekday and weekends.

|  | **Weekdays**<br>**Mean ± SD** | **Weekends**<br>**Mean ± SD** | **Pairwise**<br>**comparison** |
| --- | --- | --- | --- |
| ***Actigraphy*** |  |  |  |
| Total Time in Bed | 500.4± 41.5 | 516.9 ± 61.8 | t(70)=-2.4, $p$= .02 |
| Total Sleep Time | 447.5 ± 39.6 | 462.9 ± 57.6 | t(70)= -2.4, $p$= .02 |
| Wake After Sleep Onset | 47.5 ± 25.7 | 48.9 ± 27.4 | t(70)= .24, $p$= .48 |
| Sleep Efficiency | 89.5 ± 4.8 | 89.6 ± 4.9 | t(70)= .40, $p$= .80 |
| Sleep Onset Latency | 3.3 ±1.9 | 3.4 ± 2.7 | t(70)= -.23, $p$= .82 |
| ***App-based Phone Tracking*** |  |  |  |
| Daily Social Media Use (mins) | 185.9±123.4 | 201.6±115.3 | t(79)= -1.63, $p$= .11 |

### Cross-Sectional Analyses: Associations Between Sleep, Social Media Use, and Psychopathology

#### Data reduction

To minimize redundancy among actigraphy-derived sleep indices, a PCA was conducted on ten summary metrics from the two-week monitoring period. Four components with eigenvalues > 1 were extracted, explaining 75% of the total variance (see Supplementary Tables S4-S7 and Figure S1). Inspection of the rotated solution indicated two interpretable latent clusters representing Sleep Continuity (SleepC) and Sleep Duration Weekend Variability (SleepDWV). The SleepC factor was characterized by sleep continuity and latency indices. The SleepDWV factor was characterized by shorter average sleep duration, greater night-to-night variability in sleep duration, and larger weekday-weekend differences in sleep timing and duration. For parsimony, regression-based factor scores for the components within each cluster were averaged after aligning loadings so that higher values reflected poorer continuity or shorter and more variable sleep (see Supplementary Results for the full PCA output). These two composite indices (SleepC and SleepDWV) served as sleep predictors in subsequent models.

#### Moderation by Social Media Use

A moderation model tested whether SMU duration altered the association between Sleep Duration Weekend Variability (SleepDWV) and psychopathology symptoms (YSR Total Problems), controlling for age, sex, and Sleep Continuity (SleepC). The overall model accounted for 17% of the variance in psychopathology symptoms (R^2^=17.3, F(6, 59) = 2.22, *p* = .05). Greater SleepDWV was associated with higher psychopathology symptoms, *b* = 6.04, *SE* = 2.63, *t* = 2.30, *p* = .025, 95% CI [0.78, 11.29]. Age was a significant covariate (*b* = –1.27, *SE* = 0.62, *p* = .045) indicating older adolescents reported fewer symptoms. There was no significant effect of sex (b = –2.40, SE = 2.73, p = .38). Importantly, the SleepDWV × SMU interaction was nonsignificant, *b* = 1.79, *SE* = 2.76, *p* = .52, thus, in this sample, we did not detect moderation by SMU of the association between sleep disruption (i.e., reduced sleep duration and greater weekend-weekday sleep duration variability) and mental health problems (Figure 1A).

**Figure 1.**
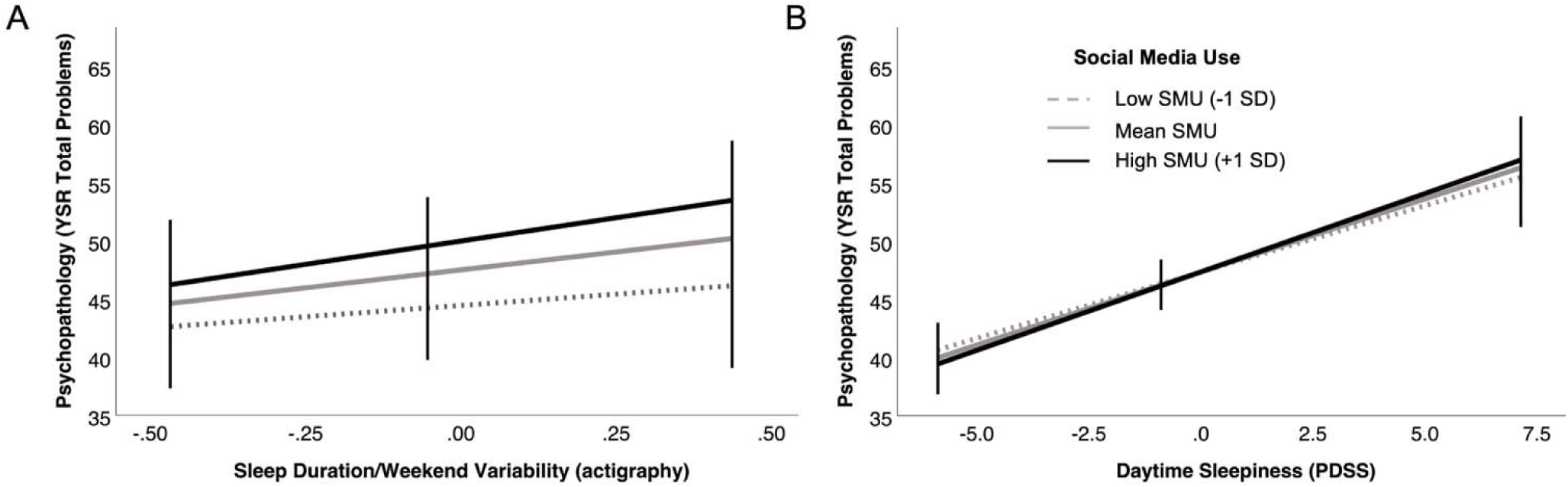
Social Media Use duration does not moderate the relationship between sleep and psychopathology symptoms. **(A)** Interaction between Sleep Duration/Weekend Variability (Sleep DWV) and social media use duration predicting psychopathology (YSR Total Problems t-scores). Predicted values are plotted at low (–1 SD), mean, and high (+1 SD) levels of standardized daily social media use (SMU) duration. Greater Sleep DWV (reflecting shorter average sleep duration and greater weekend-weekday variability in sleep duration) was associated with higher psychopathology scores across all SMU levels. The Sleep DWV × SMU interaction was nonsignificant (p = .52), indicating that the strength of the sleep–psychopathology association did not differ by social media use. All effects are adjusted for age, sex and sleep continuity metrics. **(B)** Interaction between subjective ratings of daytime sleepiness (PDSS(Drake et al., 2003) total score) and SMU duration predicting psychopathology (YSR(Achenbach, 1999) Total Problems t-scores). Predicted values are shown at low (–1 SD), mean, and high (+1 SD) levels of standardized daily SMU. Higher daytime sleepiness was associated with greater psychopathology across all levels of SMU. The PDSS × SMU interaction was nonsignificant (p = .63), indicating that the strength of the association between sleepiness and mental health did not differ by SMU duration. All effects are adjusted for age and sex.

A second moderation model replaced SleepDWV with SleepC as the predictor. The model explained 18.2% of the variance in symptoms, *F*(6, 59) = 2.17, *p* = .057. SleepC was not associated with psychopathology (*b* = 0.06, *SE* = 1.29, 95% CI [–2.52, 2.64], *p* = .96), and the SleepC × SMU interaction was not significant (*b* = 1.61, *SE* = 1.54, 95% CI [–1.47, 4.68], *p* = .30). Age showed a marginal association with symptoms (*b* = –1.20, *SE* = 0.62, 95% CI [–2.45, 0.04], *p* = .057), indicating that older adolescents tended to report fewer symptoms. Sex was not significant (*p* = .45). SleepDWV remained a significant covariate in this model (*b* = 5.90, *SE* = 2.60, 95% CI [0.71, 11.10], *p* = .027). After applying FDR correction across the two moderation models, the main effect of SleepDWV remained significant. Across models, we did not detect evidence that SMU moderated any sleep-mental health associations.

To evaluate whether the association between SleepDWV and psychopathology varied by symptom domain, separate regression models were conducted predicting internalizing and externalizing symptoms, controlling for age, sex, and SleepC. The model predicting externalizing symptoms was significant (*R²* = .12), *F*(3, 65) = 2.89, *p* = .042. Greater SleepDWV predicted higher externalizing problems (*b* = 5.72, *SE* = 2.05, 95% CI [1.62, 9.82], *p* = .007). In contrast, the model predicting internalizing symptoms was not significant (*R²* = .07), *F*(3, 65) = 1.60, *p* = .199, and the effect of SleepDWV did not reach significance (*b* = 4.92, *SE* = 2.69, 95% CI [–0.46, 10.30], *p* = .072). Age, sex, and SleepC were not significant predictors in either model (all *ps* > .15). These findings indicate that shorter and more variable sleep is significantly associated with greater externalizing symptoms, rather than internalizing symptoms.

To aid interpretation of the SleepDWV factor, an exploratory analysis examined two actigraphy-derived sleep metrics: mean total sleep time (TST) across the two-week monitoring period and variability in TST (indexed as the standard deviation of TST across monitoring days). Greater variability in sleep duration predicted higher psychopathology symptoms (b = 0.23, SE = 0.06, 95% CI [0.11, 0.36], *p* < .001), whereas mean sleep duration was not associated with symptoms (b = −0.01, SE = 0.03, 95% CI [−0.08, 0.05], p = .682). These findings suggest that the association captured by the SleepDWV factor was more closely related to variability in sleep duration than average sleep duration.

#### Daytime sleepiness

To examine whether self-reported daytime sleepiness showed the same pattern as the objective sleep metrics, we tested PDSS scores as the predictor of psychopathology. This model explained 48% of the variance in YSR Total Problems, F(5, 72) = 13.43, p < .001. Higher PDSS scores predicted more symptoms (b = 1.25, SE = 0.16, t = 7.59, p < .001, 95% CI: 0.92, 1.58). SMU did not moderate this relationship (interaction b = 0.09, SE = 0.19, p = .63; Figure 1B).

#### Direct effects of social media use

To assess whether SMU duration showed any direct associations with sleep or mental health, we examined Pearson’s correlations between SMU duration and all composite sleep and symptom variables. No significant associations were observed (all *p*s > .10, *r*s = .03–.20). This indicates that the absence of moderation is not masking an underlying direct effect, i.e. SMU duration was not significantly associated with sleep or psychopathology.

*Daily Within-Subjects Associations Between Sleep, Affect, Mood and Social Media Use* Linear mixed-effects models tested day-to-day within-person associations among sleep, affect, and SMU durations. The models simultaneously estimated within-person daily associations and between-person differences across participants. None of the within- or between-person associations described below remained significant following FDR correction. Unadjusted estimates are reported to characterize the direction and magnitude of associations across the daily models. At the unadjusted level, nights with longer than usual total sleep time predicted higher positive affect the following day (b = 0.0026 ± 0.0011, t(738.1) = 2.40, *p* = .017) and lower negative affect (b = - 0.004±0,0016, t(741.0)= -2.38, *p* = .018). Shorter-than-usual sleep predicted more time spent on social media the next day (b = –0.073 ± 0.036, t (749.7)= -2.02, *p* = .043). No reverse associations emerged (i.e., daily SMU did *not* predict same-night sleep or next-day affect). With respect to age and sex, older adolescents (b = 13.11 ± 4.96, t(70.5) = 2.64, *p* = .01) and boys (b = 55.66 ± 22.08, t(69.7) = 2.52, *p* = .014) spent more time on social media overall. Additional between-subjects estimates indicated that, at the unadjusted level, adolescents with greater average SMU showed shorter mean sleep duration across the monitoring period (b = −0.103 ± 0.047, t(66.7) = −2.17, *p* = .033). Greater average mood elation was also associated with higher average SMU duration (b = 1.38 ± 0.64, t(79.3) = 2.16, *p* = .034).

## DISCUSSION

This study examined the relationships between sleep, social media use (SMU), and mental health concurrently in adolescents using a multimodal assessment approach: wrist actigraphy to capture sleep, smartphone screen-time logs to quantify SMU duration and daily ecological momentary assessments (EMA) of mood and affect. By continuously monitoring adolescents for two weeks during the school term, we captured habitual sleep patterns, weekday-weekend variability in sleep, daily affect, and objective SMU duration. Across the cross-sectional analyses, disrupted sleep characterized by shorter sleep duration and greater variability of sleep duration from weekdays to weekends was associated with higher psychopathology symptoms, particularly externalizing problems. Follow-up analyses indicated that the cross-sectional association was more closely related to variability in sleep duration than average sleep duration, suggesting that irregular sleep schedules may be particularly relevant to adolescent psychopathology. In contrast, the amount of time adolescents spent on social media did not predict psychopathology nor did it alter the relationship between sleep and mental health. These findings suggest that variability in sleep, rather than SMU duration, is a more robust correlate of adolescents’ emotional and behavioral functioning.

Daily tracking of sleep, affect, mood, and SMU over two weeks provided an opportunity to examine temporal dynamics within individuals. Before correction for multiple comparisons, nights when adolescents slept longer than their own average were followed by lower negative affect and higher positive affect the next day, and shorter-than-usual sleep predicted greater time spent on social media the following day. Notably, we did not observe the reverse pattern; daily SMU did not predict sleep, affect or mood stability. Although these effects did not survive corrections, the consistent directionality across models suggests that sleep may show stronger temporal associations with affect than daily variation in SMU duration. Across participants, higher average SMU was associated with shorter average sleep duration and greater mood elation across the monitoring period, although these effects likewise did not survive correction for multiple comparisons. These findings align well with prior work showing that poorer-than-usual subjective sleep predicts heightened negative affect in young adolescents, especially those with higher externalizing symptoms(Evans et al., 2024). Similarly, another intensive longitudinal study with actigraphy found that better nightly sleep predicts modest increases in next-day positive affect, whereas high arousal during the day may shorten same-night sleep (Shen et al., 2022). Similar short-term effects have been observed in adults, where brief two-to-four-day monitoring periods show that subjective poor sleep predicts higher next-day anxiety(Ben Simon et al., 2020). Converging with these EMA findings, our prior work demonstrated that oscillatory activity during non-rapid eye movement sleep is associated with reduced negative affect and anxiety upon awakening(Arpaci et al., 2024), collectively underscoring sleep’s central role in regulating emotional processes across temporal and physiological levels of analysis. Taken together, these findings suggest that sleep may be more consistently associated with affective functioning than duration-based measures of social media use. Although the present observational study cannot determine whether improving sleep would reduce psychopathology symptoms, emerging intervention studies suggest that sleep may represent a promising and potentially modifiable target for youth mental health and well-being (Dewald-Kaufmann et al., 2014, Van Dyk et al., 2017, Blake et al., 2018). Additional intervention research is needed to determine whether the associations observed here reflect causal effects and to identify which aspects of sleep are most relevant for adolescent mental health.

In contrast to the robust associations observed for sleep, no parallel effects were identified for social media use. The lack of direct associations between SMU duration and mental health suggests that duration-based indices of online engagement may be relatively weak predictors of youth well-being. We also did not detect moderation by SMU of the sleep-psychopathology relationship, although this finding should be interpreted cautiously given the sample size available for between-subject interaction analyses. These findings are broadly consistent with prior EMA research in young adults showing that self-reported bedtime SMU was not associated with poorer sleep or next-day affective wellbeing (Das-Friebel et al., 2020). Our findings extend this work to adolescents during the school term and suggest that objectively measured SMU duration is likewise a weak predictor of daily sleep and affective functioning. In fact, meta-analytic evidence reveals small correlations observed between overall SMU and mental health, whereas moderate effects emerge for problematic use(Ahmed et al., 2024). However, in most studies, problematic SMU is assessed through self-report scales (e.g. the Bergen Facebook Addiction Scale(Andreassen et al., 2012) modeled after behavioral addiction frameworks and capturing symptoms such as preoccupation, tolerance, withdrawal, and loss of control(Huang, 2022). These instruments may overlap with mental health measures, raising concerns that observed associations reflect measurement rather than distinct causal processes. Objective smartphone-based metrics, such as those employed here, help circumvent these shared-method biases and provide a stronger test of whether time-based exposure alone predicts mental health outcomes. Importantly, the absence of associations may also reflect a very small effect of SMU on sleep or mental health at the population level(Bauducco et al., 2024). Future research should move beyond duration-based indices to examine the content, context, and emotional tone of digital engagement (e.g., active vs. passive use, interpersonal interactions, emotional valence of posts, timing relative to sleep) as well as disparities in technology access and sleep environments across diverse adolescent populations with the dual goals of characterizing population-level effects and identifying subgroups of adolescents who may be particularly vulnerable to adverse effects of digital engagement.

Several limitations should be noted. Although sleep was measured objectively using actigraphy, which strengthens confidence in the validity of daily sleep estimates, the nature and size of the current sample may have limited power to detect smaller effects in both within- and between-subjects analyses, particularly smaller interaction effects in the between-subjects moderation models. Future studies should include larger samples enriched for psychopathology to better capture variability across different psychopathology dimensions. Second, the association between daytime sleepiness and psychopathology should be interpreted in light of limited measurement overlap, as the YSR Total Problems scale includes a single sleepiness item. Although this item may contribute to the association, it is unlikely to account for the overall magnitude of the relationship. Further, the sample was predominantly White, with mostly college-educated parents, and families characterized by low household chaos and high environmental structure, which may limit generalizability. Adolescents from more socioeconomically or racially diverse backgrounds may experience different sleep-social media-mental health dynamics due to social stressors, resource disparities, or environmental instability. Social determinants such as access to technology, housing conditions, and disparities in sleep environments could influence social media exposure and sleep in ways not reflected here. Further, although objective smartphone-based SMU tracking was employed, the proprietary algorithms used by different operating systems limit the transparency regarding how SMU was classified and quantified, and differences across devices may have introduced additional variability in SMU estimates. SMU on secondary devices also could not be objectively quantified. Additionally, our objective SMU measure quantified daily duration but did not capture the timing of SMU across the day. This is an important limitation because nighttime engagement is more likely to displace sleep opportunity. Future work should develop measures that capture not only how much adolescents engage with social media, but also when they engage and what they encounter. Examining interactions between timing and content may be particularly important for identifying the aspects of digital media use most relevant to sleep and mental health. Finally, the two-week observation window may not have been sufficient to establish a stable baseline of adolescents’ habitual sleep patterns, capture the full range of day-to-day fluctuations or reliably examine differences between weekdays and weekends. Longer monitoring periods may reveal stronger longitudinal daily associations between sleep and affect that were not detectable in the current dataset.

Overall, the present study suggests that sleep may be more consistently linked to adolescent mental health than duration-based measures of social media use. The results also suggest that duration-based measures of social media use may provide limited insight into adolescent mental health when considered in isolation. Instead, prevention and intervention efforts should consider the broader ecosystem of adolescent functioning, including adequate and consistent sleep schedules and the quality (not just the quantity) of digital engagement. From a policy standpoint, the present findings support approaches that consider both sleep and the broader context of adolescents’ digital experiences, rather than focusing exclusively on time-based restrictions. Future research should continue to integrate objective, high-frequency measures of sleep and digital behavior to inform interventions that address the complex and reciprocal relationships between technology use, sleep, and emotional well-being in youth.

## Funding Information and Acknowledgments

This work was supported by a University of Iowa Scanlan Center for School Mental Health Research Grant to BB. The authors declare no competing or potential conflicts of interest.

## Data availability statement

The data that support the findings of this study are available from the corresponding author upon reasonable request.

## Ethics Statement

Written informed parental consent and assent from participants younger than 18 years of age and written informed consent from participants aged 18 years was appropriately obtained. The study protocol was approved by the University of Iowa Institutional Review Board on 9/11/2023, protocol number IRB-01 202306574.

## Abbreviations

SMU: Social Media Use
EMA: ecological momentary assessments

