## Supplementary Information for "When “Time Online” Tells Us Little: Duration of Social Media Use does not Influence Sleep Disruption or Psychopathology in Adolescents"

**Supplementary Methods**

**Table S1.** Morning EMA Survey

| Question | Response |
| --- | --- |
| 1. What time did you go to bed last night? | Open response |
| 2. How long did it take you to fall asleep after lights out? | Open response |
| 3. Where did you sleep (e.g. own bed, couch, friend’s house)? | Open response |
| 4. What time did you wake up this morning? | Open response |

**Table S2.** Evening EMA Survey

| Question | | Response |
| --- | --- | --- |
| Instructions: The following questions consist of a number of words that describe different feelings and emotions. Read each item, then indicate to what extent you have felt that way today using the provided scale. | | |
| 1. Cheerful | 1= very slightly or not at all, 2= a little, 3= moderately, 4= quite a bit, 5=extremely | |
| 2. Relaxed | 1= very slightly or not at all, 2= a little, 3= moderately, 4= quite a bit, 5=extremely | |
| 3. Irritable | 1= very slightly or not at all, 2= a little, 3= moderately, 4= quite a bit, 5=extremely | |
| 4. Sad | 1= very slightly or not at all, 2= a little, 3= moderately, 4= quite a bit, 5=extremely | |
| 5. Calm | 1= very slightly or not at all, 2= a little, 3= moderately, 4= quite a bit, 5=extremely | |
| 6. Afraid | 1= very slightly or not at all, 2= a little, 3= moderately, 4= quite a bit, 5=extremely | |
| 7. Happy | 1= very slightly or not at all, 2= a little, 3= moderately, 4= quite a bit, 5=extremely | |
| 8. Guilty | 1= very slightly or not at all, 2= a little, 3= moderately, 4= quite a bit, 5=extremely | |
| 9. Nervous | 1= very slightly or not at all, 2= a little, 3= moderately, 4= quite a bit, 5=extremely | |
| 10. Lonely | 1= very slightly or not at all, 2= a little, 3= moderately, 4= quite a bit, 5=extremely | |
| 11. At ease | 1= very slightly or not at all, 2= a little, 3= moderately, 4= quite a bit, 5=extremely | |
| 12. Enthusiastic | 1= very slightly or not at all, 2= a little, 3= moderately, 4= quite a bit, 5=extremely | |
| Instructions: For the following questions, rate how much you agree with the statement. | | |
| 13. Right now I feel much more giddy, silly, or happy than usual. | | *Sliding scale from 0-100 |
| Instructions: Answer the questions below | | |
| 14. Did you take any naps today? | | YES or NO |
| 14b. IF YES to #14, what time did you start and wake up from your nap? | | Open response |
| 15. Did you take the watch off today? | | YES or NO |
| 16b. IF YES to #16, what times did you take it off and put it back on? | | Open response |

**Supplementary Results**

**Table S3.** Research Diagnoses based on the Kiddie Schedule for Affective Disorders and Schizophrenia (KSADS)

| 1 | Social Anxiety Disorder |
| --- | --- |
| 2 | Obsessive Compulsive Disorder |
| 3 | Generalized Anxiety Disorder |
| 4 | Obsessive Compulsive Disorder |
| 5 | Social Anxiety Disorder |
| 6 | Major Depressive Disorder, Obsessive Compulsive Disorder |
| 7 | Major Depressive Disorder, Suicidality |
| 8 | Obsessive Compulsive Disorder |
| 9 | Major Depressive Disorder |
| 10 | Adjustment Disorder with Anxiety |
| 11 | Major Depressive Disorder, Alcohol Use Disorder, Drug Use Disorder |
| 12 | Major Depressive Disorder |
| 13 | Major Depressive Disorder, Generalized Anxiety Disorder, Obsessive-Compulsive Disorder |
| 14 | Major Depressive Disorder |
| 15 | Major Depressive Disorder, Obsessive Compulsive Disorder |

**Principle Components Analysis (PCA) of actigraphy measured sleep:**

To reduce redundancy among the 14-day actigraphy-derived sleep measures, a principal component analysis (PCA) was performed on 10 indices: means and standard deviations (SD) of Total Sleep Time (TST), Sleep Onset Latency (SOL), Sleep Efficiency (SE), and Wake After Sleep Onset (WASO) across 14 days, along with two measures of weekday-weekend sleep misalignment, Social Jetlag (weekday-weekend discrepancy in sleep duration) and Sleep Debt (mid-sleep difference between weekdays and weekends).

All variables were entered into the PCA using the correlation matrix with Varimax rotation. Extraction communalities indicated generally good representation of the variables by the retained components (e.g., mean TST = .81; mean SOL = .90; mean SE = .93; mean WASO = .89; SD SE = .88; SD WASO = .90; SD SOL = .88; SD TST = .54; Social Jetlag = .46; Sleep Debt = .32). Using the eigenvalue > 1 criterion, four components were initially retained, accounting for 75.02% of total variance prior to rotation (Component 1: eigenvalue = 3.08, 30.83%; Component 2: 1.88, 18.83%; Component 3: 1.51, 15.10%; Component 4: 1.03, 10.26%). After Varimax rotation, the cumulative percentage of variance explained by the four components was 28.13%, 47.15%, 62.29%, and 75.02%, respectively.

- Component 1 was characterized by high loadings from SD WASO (.93), SD SE (.87), mean WASO (.76), and inverse mean SE (–.74) and was interpreted as the measure of “*Sleep Continuity*”.
- Component 2 included mean SOL (.95) and SD SOL (.94) and was interpreted as the measure of “*Sleep Latency*”.
- Component 3 included SD TST (.70), Social Jetlag (.66), inverse Sleep Debt (–.46) and mean WASO (-.33) was interpreted as a measure of “*Weekend Variability in Sleep Duration and Timing*”.
- Component 4 included mean TST (.87) with cross-loadings from mean SE (.51) and mean WASO (–.42) and was interpreted as a measure of “*Sleep Duration*”.

Although these four components met the eigenvalue > 1 criterion, inspection of the rotated solution revealed conceptual overlap forming two broader clusters: (1) a Sleep Continuity cluster (Components 1 + 2’ “**Sleep C**”) and (2) a Sleep Duration Weekend Variability cluster (Components 3 + 4, “**Sleep DWV**”).

For parsimony, regression-based factor scores were computed for all four rotated components and then averaged within each cluster, after sign-harmonizing loadings so that higher values reflected poorer sleep continuity or shorter and more variable sleep duration. The resulting two composite scores were re-standardized (mean = 0, SD = 1) for use in subsequent moderation models.

**Table S4.** Correlation matrix of the 10 actigraphy-derived sleep variables used in the PCA.

| **Variables** | mean TST | mean SOL | mean SE | mean WASO | SD TST | SD SOL | SD SE | SD WASO | Social Jetlag | Sleep Debt |
| --- | --- | --- | --- | --- | --- | --- | --- | --- | --- | --- |
| **mean TST** | 1.00 | .075 | .353 | -.246 | -.141 | .021 | -.242 | -.090 | -.057 | .099 |
| **mean SOL** | .075 | 1.00 | -.211 | .151 | .037 | .803 | .015 | -.067 | -.018 | .084 |
| **mean SE** | .353 | -.211 | 1.00 | -.986 | .035 | -.126 | -.488 | -.515 | .061 | -.040 |
| **mean WASO** | -.246 | .151 | -.986 | 1.00 | -.083 | .071 | .463 | .543 | -.077 | .051 |
| **SD TST** | -.141 | .037 | .035 | -.083 | 1.00 | .033 | .276 | .107 | .249 | -.047 |
| **SD SOL** | .021 | .803 | -.126 | .071 | .033 | 1.00 | .051 | -.094 | .103 | .045 |
| **SD SE** | -.242 | .015 | -.488 | .463 | .276 | .051 | 1.00 | .895 | .116 | -.234 |
| **SD WASO** | -.090 | -.067 | -.515 | .543 | .107 | -.094 | .895 | 1.00 | .054 | -.151 |
| **Social Jetlag** | -.057 | -.018 | .061 | -.077 | .249 | .103 | .116 | .054 | 1.00 | .131 |
| **Sleep Debt** | .099 | .084 | -.040 | .051 | -.047 | .045 | -.234 | -.151 | .131 | 1.00 |

*TST* = total sleep time; *SOL* = sleep onset latency; *SE* = sleep efficiency; *WASO* = wake after sleep onset. Values represent Pearson correlations across participants averaged over 14 days of actigraphy monitoring.

**Table S5.** Extraction communalities.

| Variable | Extraction Communality |
| --- | --- |
| mean TST | 0.81 |
| mean SOL | 0.90 |
| mean SE | 0.93 |
| mean WASO | 0.89 |
| SD TST | 0.54 |
| SD SOL | 0.88 |
| SD SE | 0.88 |
| SD WASO | 0.90 |
| Social Jetlag | 0.46 |
| Sleep Debt | 0.32 |

**Table S6.** Eigenvalues before and after Varimax rotation.

| Component | Eigenvalue (Initial) | % Variance Explained (Initial) | Eigenvalue (Rotated) | % Variance Explained (Rotated) |
| --- | --- | --- | --- | --- |
| Component 1 | 3.083 | 30.832 | 2.813 | 28.126 |
| Component 2 | 1.883 | 18.828 | 1.903 | 19.027 |
| Component 3 | 1.510 | 15.097 | 1.514 | 15.135 |
| Component 4 | 1.026 | 10.260 | 1.273 | 12.729 |

**Table S7**. Rotated component matrix showing standardized loadings for all 10 variables.

| Variable | Component 1 (Sleep Continuity) | Component 2 (Sleep Latency) | Component 3 (Weekend Variability) | Component 4 (Sleep Duration) |
| --- | --- | --- | --- | --- |
| mean TST |  |  |  | .865 |
| mean SOL |  | .945 |  |  |
| mean SE | -.742 |  |  | .508 |
| mean WASO | .763 |  | -.328 | -.419 |
| SD TST |  |  | .704 |  |
| SD SOL |  | .936 |  |  |
| SD SE | .871 |  |  |  |
| SD WASO | .927 |  |  |  |
| Social Jetlag |  |  | .664 |  |
| Sleep Debt |  |  | -.455 |  |

**Table S8.** Within-subject correlations among daily sleep, affect, mood elation, and social media use variables. Correlations were calculated using person-mean centered observations and reflect day-to-day associations within individuals. Values shown are Pearson correlation coefficients.

| **Variable** | **1** | **2** | **3** | **4** | **5** | **6** |
| --- | --- | --- | --- | --- | --- | --- |
| 1. Sleep TST | 1 |  |  |  |  |  |
| 2. Sleep WASO | 0.030 | 1 |  |  |  |  |
| 3. Phone SMU | -0.069* | 0.004 | 1 |  |  |  |
| 4. Positive Affect^a^ | 0.002 | 0.001 | -0.085* | 1 |  |  |
| 5. Negative Affect^a^ | 0.008 | -0.049 | 0.046 | -0.338* | 1 |  |
| 6. Mood Elation | -0.031 | 0.027 | -0.067* | 0.321* | -0.219* | 1 |

TST = Total Sleep Time; WASO = Wake After Sleep Onset; SMU = Social Media Use; **p* < .05

^a^based on a modified 12-item version (Shen et al., 2022) of the Positive and Negative Affect Schedule (PANAS-X (Watson, 1999).

**Table S9.** Between-subject correlations among sleep, affect, mood elation, and social media use variables. Correlations were calculated using participant-level means across the monitoring period and reflect average differences between participants across the monitoring period. Values shown are Pearson correlation coefficients.

| **Variable** | **1** | **2** | **3** | **4** | **5** | **6** |
| --- | --- | --- | --- | --- | --- | --- |
| 1. Sleep TST | 1 |  |  |  |  |  |
| 2. Sleep WASO | -0.242* | 1 |  |  |  |  |
| 3. Phone SMU | -0.273* | -0.125 | 1 |  |  |  |
| 4. Positive Affect^a^ | 0.025 | 0.040 | 0.141 | 1 |  |  |
| 5. Negative Affect^a^ | -0.009 | 0.098 | 0.143 | -0.214* | 1 |  |
| 6. Mood Elation | -0.108 | -0.125 | 0.235 | 0.017 | 0.264* | 1 |

TST = Total Sleep Time; WASO = Wake After Sleep Onset; SMU = Social Media Use; **p* < .05

^a^based on a modified 12-item version (Shen et al., 2022) of the Positive and Negative Affect Schedule (PANAS-X (Watson, 1999).

**Figure S1.** Scree plot of eigenvalues from the principal components analysis (PCA) of actigraphy-derived sleep variables. Four components had eigenvalues greater than 1, explaining 75% of total variance. The inflection in the curve and conceptual overlap among the components supported a solution with two interpretable latent clusters representing Sleep Continuity and Sleep Duration Weekend Variability.

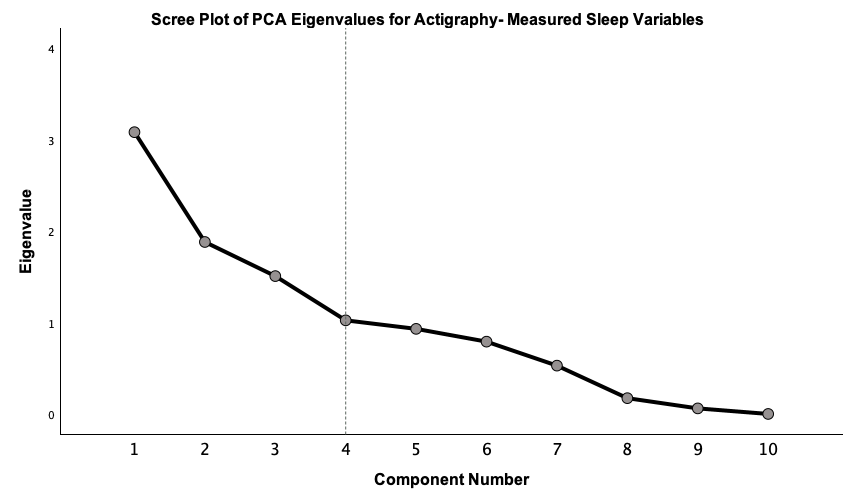
